# The salivary microbiome shows a high prevalence of core bacterial members yet variability across human populations

**DOI:** 10.1101/2021.12.13.471511

**Authors:** Xinwei Ruan, Jiaqiang Luo, Pangzhen Zhang, Kate Howell

**Affiliations:** School of Agriculture and Food, Faculty of Veterinary and Agricultural Sciences, University of Melbourne, Parkville 3010, Australia

**Keywords:** Microbiome, Oral microbiome, Westernization, Dietary pattern

## Abstract

Human saliva contains diverse bacterial communities, reflecting human health status, dietary patterns and contributing to variability in the sensory perception of food. Many descriptions of salivary microbiome diversity compare commonalities and differences with reference to a diseased state, but the composition of healthy saliva has not been described. Here, we use a meta-analysis approach to define and explore the core membership of the human salivary microbial community by collecting and re-analysing raw 16S rRNA amplicon sequencing data from 47 studies with 2206 saliva samples. We found 68 core bacterial taxa that were consistently detected. Differences induced by various host intrinsic and behaviour factors, including gender, age, geographic location, tobacco usage, and alcohol consumption, were evident. The core of the salivary microbiome was verified by collecting and analysing saliva in an independent study. These results suggest that the methods used can effectively define a core microbial community in human saliva with high prevalence. The core salivary microbiome demonstrated both stability and variability among populations. Geographic location was identified as the host factor with the largest effect in shaping salivary microbiota. The independent analysis underlined that the impact of geographic variation is likely due to diet.

## Introduction

Human saliva plays an essential role in influencing the sensory perception of foods and beverages and driving the purchase decisions of consumers. When food is taken into the mouth, mixing and mastication allow a semi-solid bolus to be formed, and at the same time, aroma and flavour to be released from the food [1]. The variation in perceived responses can be attributed to the inter-individual variability in salivary composition, especially salivary microorganisms. As a complex ecosystem, the human oral cavity hosts thousands of bacterial taxa, interacting with themselves, other microorganisms and the broader cavity [2]. It is an ecological system that contains many distinct sub-niches, including saliva, dental plaques, gingival sulcus, epithelial cells on the cheek, tongue, and teeth [3]. High heterogeneity has been reported between the composition of microbial communities that colonise on different sites [4]. Saliva is recognised as a reservoir of microorganisms from all ecological niches in the human mouth with long-term stability [5]. The ensemble of microorganisms and the expressed genetic material in human saliva; the ‘salivary microbiome’ is crucial to understanding the healthy state of humans, and indeed, what this means for sensory appreciation of foods and beverages.

The contribution of salivary microbiome to sensory perception of foods has been described by various studies [6-8]. However, the diverse conclusions suggest that the role of the salivary microbiome may be confounded by inter-individual variance. Characterising the microbial communities commonly found in most human saliva regardless of the study-specific variation could help establish the connection between salivary composition and food preference. Comparing results from different studies introduces significant technical and bioinformatic biases [9] especially when studies have targeted different 16S rRNA hypervariable regions for amplification [10]. On this basis, the shifts from a common salivary microbiome by diseases or host lifestyle factors will also be more prominent. A meta-analysis of both sequencing data and associated meta-data can summarise the existing knowledge and identify the commonalities and differences in salivary microbiota between people from various backgrounds.

Attempts to define the core oral microbiome of a healthy human has occurred during the last ten years [11]. The core microbiome is described as the common group of microbes that are important for host biological function [12]. It provides a foundation for prioritising members adapted to the host environment [13]. Although the shift in human salivary composition caused by diseases has been studied for decades [14-19], our understanding of the impact of host intrinsic and behaviour factors is still limited.

Many host characteristics have been shown to have an impact on the composition of the salivary microorganisms, including age [20], diet [21, 22], ethnicity [23], gender [24], smoking [25], alcohol use [26], circadian rhythm [27], body mass index [28], and the type of stimulation [29]. Some studies have correlated the diverse microbiome with the distinct sensory responses between consumer groups [30, 31] It has also been reported that people from different countries are colonised with distinct salivary bacterial communities [32]. Li et al. analysed the human oral microbiome from Africa, Alaska, and Germany and reported differences between the human groups living in various climate conditions [33]. However, a clear global pattern in salivary microbial composition is lacking, which considers both core microbial populations and variability amongst different groups.

In this paper, we collected raw 16S rRNA sequences of human salivary microbiota from 47 publicly available datasets spanning 15 different countries. These raw data were systematically re-analysed and pooled together to define a core salivary microbiome. We classified all sequences into operational taxonomic units (OTUs) at 97% identity against the Human Oral Microbiome Database (HOMD) to minimise the technical variation induced by comparing data from different hypervariable regions. It allowed us to make a comparison between studies and reduced the redundancy in the dataset for defining the “core” microbial members. Using the metadata acquired with raw data, we also investigated the influences of several host factors and technical factors on human salivary microbiota. Factors that showed a potentially strong impact on shaping microbial communities in saliva were selected, and the taxa as potential biomarkers were identified, and linked with functional predictions. Finally, verification of the meta-analysis was sought through a targeted sampling effort. Here, saliva samples were collected from 30 healthy individuals and analysed for microbial composition to compare against the results found from the global dataset. These data confirmed the composition of the core microbiome members, but the verification of ethnic origin was not possible. Our study contributes to fundamental understandings of the stable and differential salivary microbiome across healthy adult populations. We have identified bacteria linked to particular identities of participants and points to salivary microbiome composition being linked to diet, rather than ethnic origin.

## Materials and methods

### 1. Literature search and data collection

To acquire sufficient data from healthy human saliva, available public studies related to human salivary microbiota were systematically reviewed. A literature search was performed using the combination of relative terms in EMBASE, MEDLINE and Web of Science for the studies published before November 2020 using the terms described in supplementary data (Table S1). A supplementary dataset search in NCBI’s Sequence Read Archive (SRA) was also performed using the search term ‘salivary microbiome’. The resulting studies were screened according to the criteria described in supplementary material. Raw sequence data acquired from the healthy individuals of selected studies were downloaded from SRA and European Nucleotide Archive (ENA), using SRA Toolkit.

The files were converted to the FASTQ formats if necessary. Sequence data from each selected study were processed separately using QIIME2 (version.2020.2) [34]. The resulting OTU table was exported into BIOM format. Further analyses were carried out in R (version 4.1.0) with custom scripts as detailed in supplementary material.

### 2. Defining the core microbiome

The core microbiome was determined based on the abundance-occupancy pattern. Two methods adapted from Shade and Stopnisek [35] and Wu et al. [13] were used. For the methods adapted from the study of Shade and Stopnisek [35], samples were rarefied to 5 000. Both Bray-Curtis similarity and weighted Unifrac distance were used to determine the contribution in the percentage of the prospective core set to the overall beta diversity. For the method adapted from Wu et al. [13], OTUs were filtered out with the mean relative abundance (MRA) bigger than 0.1% and the presence in more than 75% of samples or 100% occupancy in more than 10 studies. To investigate the bacteria-bacteria interactions in salivary microbial communities, the co-occurrence network was constructed using pairwise Spearman’s correlation based on relative abundance. The Spearman’s correlation was calculated using the *rcorr* function in the *Hmisc* R package [36] and visualised by Cytoscape v3.8.2 [37]. A correlation with Spearman’s correlation coefficient > 0.5 or < -0.5 and p-value < 0.01 is considered as statistically robust and shown in the network.

### 3. Functional prediction

Microbial metagenomes were inferred from 16S rRNA gene-based bacterial profiles, and the functional prediction were conducted based on Kyoto Encyclopedia of Gene and Genomes (KEGG) database [38] using the default pipeline in Phylogenetic Investigation of Communities by Reconstruction of Unobserved States 2 (PICRUSt2) (Douglas et al., 2020). The ANCOM-BC analysis was used to identify the differential abundant KEGG pathways by geographic location, adjusting for hypervariable regions. At the same time, a random forest model was established for distinguishing Chinese and Western samples, and the importance of pathways was measured using mean decreased accuracy. Spearman’s correlation was performed to assess the relationship between the relative abundance of differential pathways and genera. The significant correlations were visualised using the *corrplot* package [39] in R.

### 4. Comparison between Chinese and Western people on an independent cohort

Saliva samples were collected from 26 participants (aged 20-60 years) recruited for a wine assessment experiment and consisted of 13 Chinese and 13 Western wine experts (Table 1). The study was approved by the Office for Research Ethics and Integrity of the University of Melbourne (Ethics ID: 1852616). Each group had six female panellists and seven male panellists. The Western panellists were defined as people who have lived in Australia for more than ten years. Chinese panellists were defined as people who were born in China and had lived in Australia for no more than 18 months.

Bacteria genomic DNA was extracted from human saliva using QIAGEN® MagAttract® PowerSoil® DNA KF Kit [40] and subjected to 16S rRNA amplicon sequencing on the Illumina platform following the Earth Microbiome Project protocols (https://earthmicrobiome.org/protocols-and-standards/16s/). The raw data are available in NCBI Sequence Read Archive, with accession number PRJNA786805.

The raw sequences were processed using the same pipeline in the meta-analysis as described in section 2. Additionally, raw sequencing reads were denoised into zero-radios OTUs (zOTUs) by UNOISE3 pipeline [41], and taxonomically classified by classifiers trained on the full-length 16S rRNA gene SILVA v138 [42] database and eHOMD v15.22 [43], respectively. The affiliation between each ZOTUs and the originating OTU was determined using a customised code adapted from Stopnisek and Shade [44] and available at https://github.com/XINWEIR/SalivaryMicrobiome_MetaAnalysis. The relative abundance of taxa at the genus level in this cohort was used as the test set for the random forest model trained using the genus-level OTU assignment information in the meta-analysis.

## Results

### 1. Inclusion of studies and sequences

In this study, we extracted the 16S rRNA gene amplicon sequencing data of healthy human saliva from 47 studies (Figure 1A) [15, 16,48, 67-110]. We abstracted data from subjects who had no diagnosed disease state, hereafter named “healthy”. Of course, subjects could have had subclinical diseases or may have altered health status for undisclosed reasons, but we considered that this would be true of the wider human population and therefore able to be included in our study. A total of 107 005 868 high quality 16S rRNA sequences were obtained. After removing all samples below 2,000 reads, 2206 samples with 909 features were retained. The retained samples included studies from 15 countries (Table S2). Most studies were conducted in three geographic regions: North America, Europe, and China (Figure 1B). Most sequences included in this meta-analysis were generated from the hypervariable region “V3-V4” and “V4” (Figure 1C).

**Figure 1.**
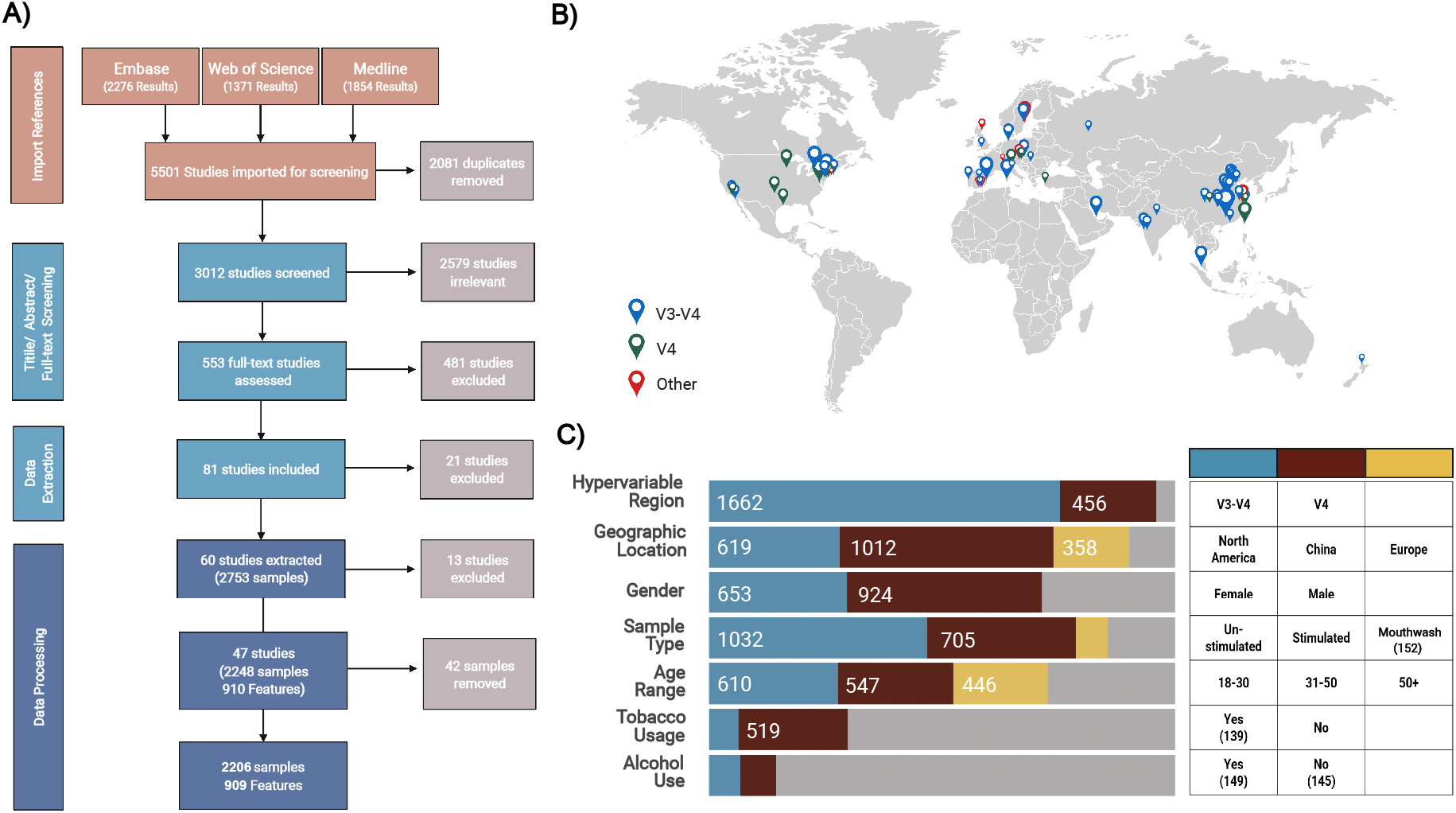
Overview of literature search procedure and metadata of included studies. **A)** Large-scale literature searching and data filtering process, followed by the number of samples submitted to the bioinformatic analyses; **B)** The locations of studies, the scale of symbols that reflect the number of samples of each study; **C)** Distribution of metadata categories.

### 2. Intrinsic and lifestyle factors have a significant effect on the host salivary microbiome

Large variability between studies was observed in the number of reads, taxonomic profile, and alpha diversity (Figure 2; Figure S1). Phylum *Bacteroidetes, Proteobacteria, Firmicutes, Fusobacteria* were dominated among all studies, while their proportion varies (Figure 2A). When studies were grouped by the geographic locations they originated from (coloured in Figure 2B, C, D), there is generally no difference between their intra-community diversity, represented by Shannon, Chao1, and Simpson indices. Only one study conducted in Qatar showed relatively lower Chao 1 index and higher Simpson’s diversity indices than studies from other locations. However, it is hard to decide whether such variation is caused by the geographic location or other technical variations.

**Figure 2.**
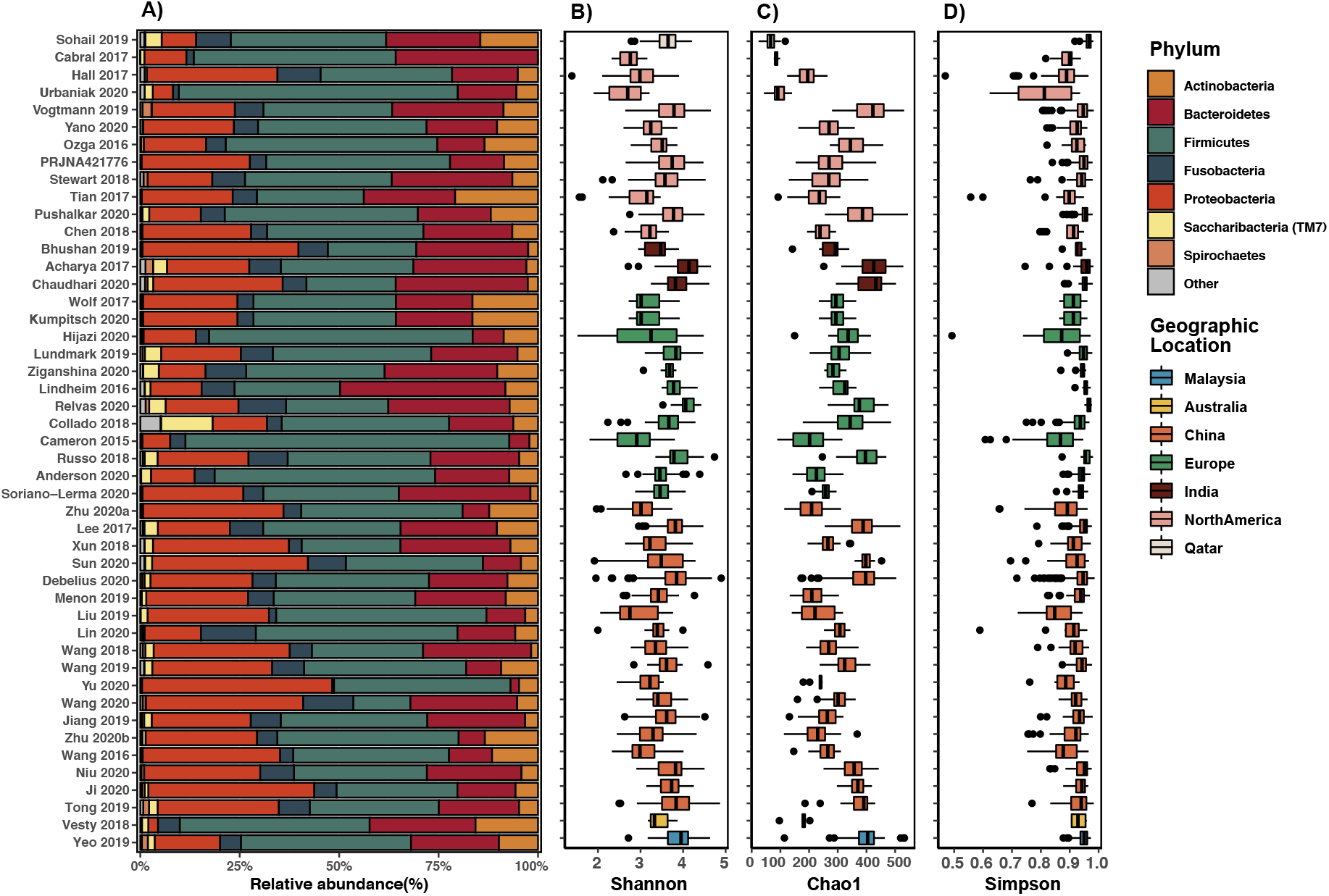
Summary of taxonomic composition and alpha diversity of included studies. **A)** The mean community composition of each study at the phylum level; The alpha-diversity measured by **B)** Shannon index; **C)** Chao 1 index; **D)** Simpson’s index, the colour of boxes stands for the geographic location of the studies. The horizontal bars within boxes represent medians. The tops and bottoms of boxes represent the 75th and 25th percentiles, respectively.

Because of the large disparity of methodologies amongst the studies used in our global analysis, we applied several different strategies for normalisation as described in Supplementary Methods. When investigating the influences of different categories using permutational multivariate analysis of variance (PERMANOVA) tests, these normalising methods were combined as appropriate with different distance metrics, including Bray-Curtis, weighted UniFrac, and Euclidean distance. Overall, the effect of rarefaction (RAR), total-sum scaling (TSS), and the rarefied relative abundance transformation (RRA) were very similar in the result of PERMANOVA (Figure 3A-D; Table S4). Meanwhile, the centred log-ratio transformation (CLR) enlarged the variance induced by an unwanted technical factor, namely the amplified hypervariable region, at all taxonomic levels. The beta-diversity analyses showed that all metadata categories measured have a significant (p<0.001) effect on the bacterial profile of human saliva at all taxonomic levels, adjusted for the study effect (Table S4). However, only limited variation among samples has been explained by these factors (R2 < 10%). In contrast, “study” accounts for around 35% of the variability between samples. At the OTU level, the combination of weighted UniFrac distance with two normalisations (TSS and RRA) best minimised the variability raised by different hypervariable regions (Figure 3D). The results of the unconstrained principal coordinate analysis (PCoA) are in agreement with the results of PERMANOVA. When using Bray-Curtis dissimilarity and Euclidean distance, the samples separated distinctly according to the hypervariable regions in PCoA plots, whereas the plot constructed using the weighted UniFrac revealed the clusters formed by the samples from different geographic locations (Figure S2). A distinct separation of samples from three main geographic locations (Figure 3F), with more than half (58.0%) of the variance explained by the first two dimensions, using weighted UniFrac distance. In contrast, the differences between locations were confounded by which hypervariable regions were sequenced in the PCoA plot for Bray-Curtis dissimilarity (Figure 3E; Figure S2D-F). The results suggest that host intrinsic and lifestyle factors significantly influence the microbial profile in human saliva, regardless of the variation induced by technical factors.

**Figure 3.**
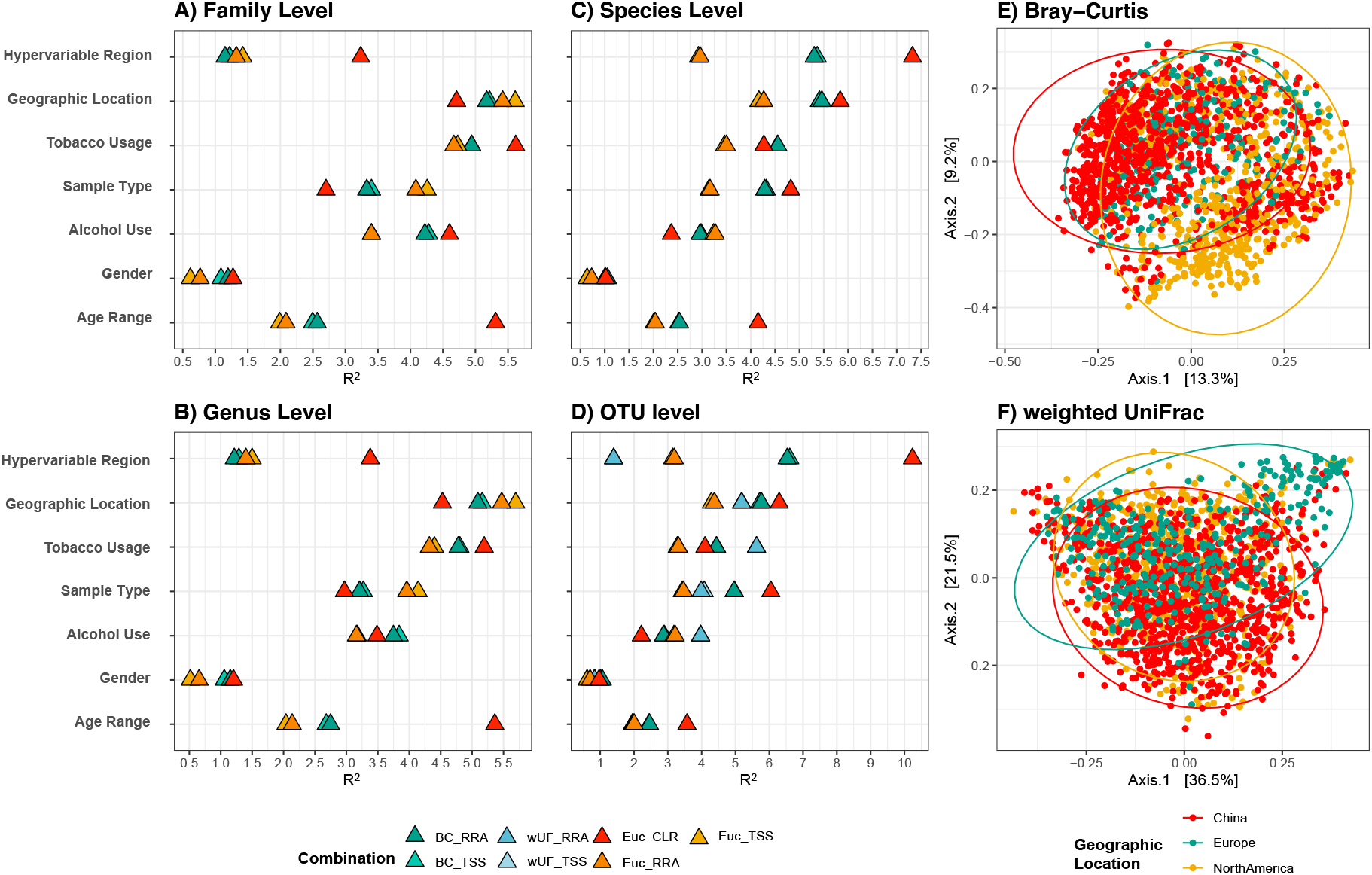
The variability in human salivary microbiota have been explained by different factors. Among them, hypervariable regions and geographic locations have the largest impact. The effect of the categories on the clustering of the sample was measured using PERMANOVA at four taxonomic levels: family **(A)**, Genus **(B)**, species **(C)** and OTU level **(D)**. The colour indicates the different combinations of normalisation (TSS, Total-sum scaling; RRA, Rarefied relative abundance; CLR, Centred log ratio) and indices (BC, Bray-Curtis; EUC, Euclidean; wUF, weighted uniFrac). Because the results of rarefication (RAR) were very close to TSS and RRA, they were not displayed in the figures. Principal coordinate analysis (PCoA) with Bray-Curtis **(E)** and weighted uniFrac **(F)** showing the differences between samples from North America, Europe, and China.

### 3. A core microbiome is defined from saliva from healthy humans

Despite the large intra- and inter-study variability, many OTUs still showed a consistently high presence and relative abundance across studies (Figure S3). These persistent OTUs detected across studies with different protocols could be functionally important for the salivary microbiome of healthy adults. We wanted to identify the most widespread microbial taxa within a specific population that allows us to better understand the broad structure of microbiomes and their potential functional consequences [12]. The abundance and occurrence frequency of taxa are two important criteria used to define the “core salivary microbiome”. Conventionally, thresholds on these two parameters filter all taxa detected, and taxa that meet both criteria can be classified as the core salivary microbiome [13]. Recently, a more standardised procedure based on abundance-occupancy distribution was proposed [35]. We employed both strategies to define the core salivary microbiome to identify the microbial features with high persistence and robustness in human saliva. Considering the sequences involved in this meta-analysis were collected from studies targeted at different hypervariable regions, close-referenced clustering at 97% identity were used to cluster sequences into OTUs. In addition, taxa defined at 100% sequence identity may increase the redundancy in the dataset [35]. Therefore, the core salivary microbiome was defined using the clustered OTUs at 97% identity. To begin, the core OTUs were determined by filtering all OTUs based on mean relative abundance and occurrence frequency using the criteria described in the Methods [13]. In total, 11.6% of all OTUs (105 OTUs) were included as Core 1 (Figure 4A: MRA + OCC; Table S5). Meanwhile, according to the method proposed by Shade and Stopnisek [35], OTUs were ranked depending on their occupancy across studies, and the contribution of top-ranked OTUs to beta-diversity was expressed by Bray-Curtis similarity and weighted UniFrac distance. Two groups of the core microbiome were prioritised by these two indices, to give different inclusions in the core, consisting of the top 69 OTUs (using weighted UniFrac; Figure 4A: BC) and 94 OTUs (using Bray-Curtis; Figure 4A: wUF) OTUs (Figure S4).

**Figure 4.**
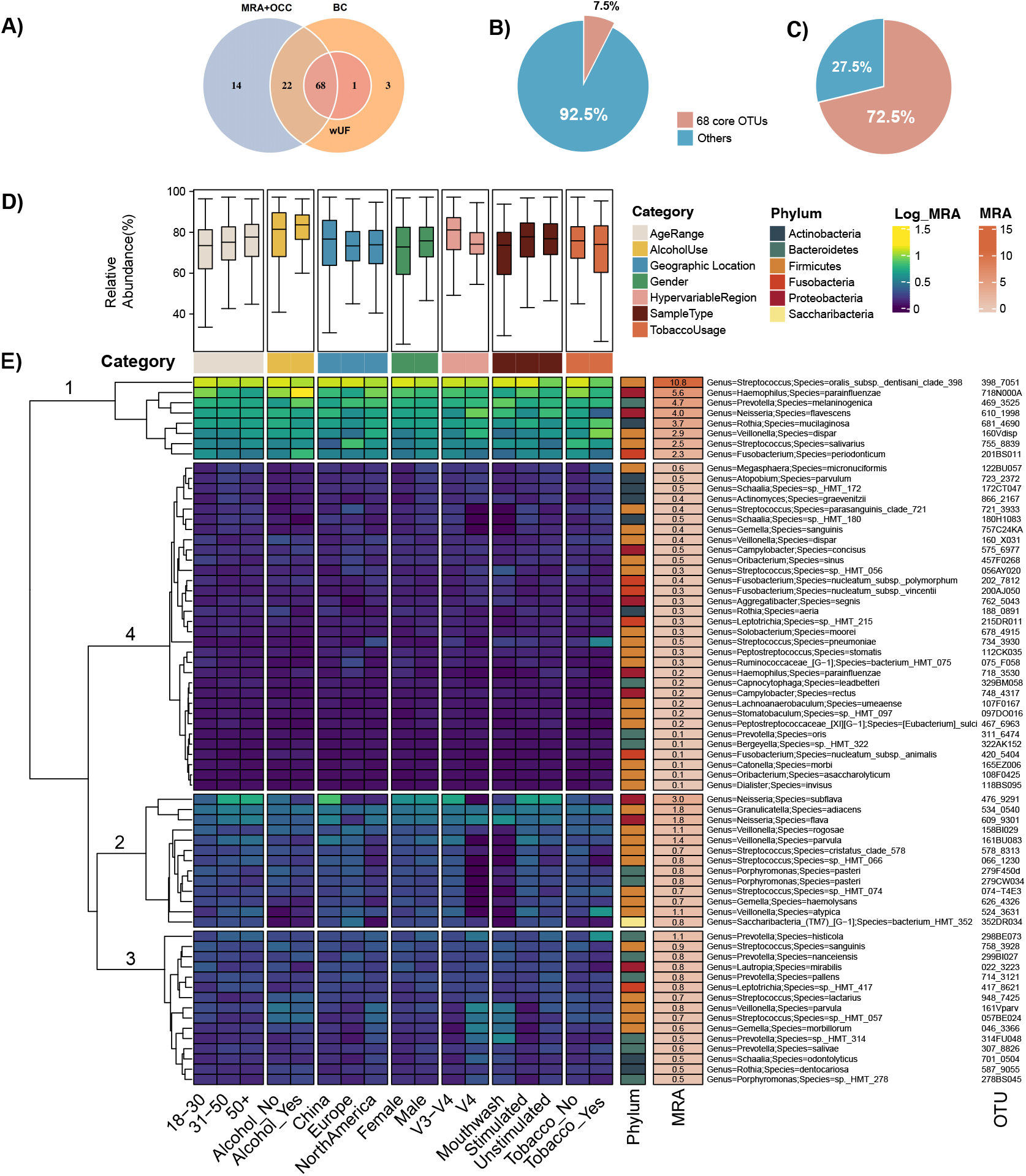
The core OTUs defined by abundance-occupancy pattern. **A)** Venn diagram showing the interaction between three methods used to define the core. Sixty-eight OTUs were defined as the core for all methods. (MRA+OCC: The thresholds were setting on mean relative abundance and occupancy to define the core; BC: The method adapted from Shade and Stopnisek using Bray-Curtis similarity; wUF: The method adapted from Shade and Stopnisek using weighted uniFrac distance). **B)** Pie chart showing the number of the core (pink) versus other OTUs (blue) identified in percentage. **C)** Pie chart showing the relative abundance of the core and other OTUs across all samples. **D)** Relative abundance of 68 core OTUs across subgroups classified by seven categories. **E)** Heatmap showing the log-transformed mean relative abundance of each core OTU at each level of different categories.

Overall, sixty-eight OTUs were shared across all three methods (Figure 4A), accounting for 7.5% of all OTUs detected and 72.5% of all 16S rRNA gene sequences after clustering and filtering (Figure 4B, C). *Firmicutes* account for nearly half (46.4 %) of all core OTUs, while only one OTU belongs to *Saccharibacteria*. The mean relative abundance (MRA) of each OTU in sub-groups classified by different factors was also measured (Figure 4D). On average, the core OTUs were highly prevalent (73.2% ± 3.4% of cumulative relative abundance) in saliva samples across different levels in subgroups classified by age, gender, geographic locations, hypervariable regions, sample type, smoking, and drinking habits. The core OTUs were clustered into four main groups based on their distribution pattern in sub-groups (Figure 4E). The eight OTUs affiliated to Cluster 1 showed overall high abundance in all sub-groups. Cluster 2 consists of core OTUs with a slightly lower mean relative abundance than Cluster 1 and higher intra-group variability. Notably, although having a higher MRA than some members of Cluster 1, “476_9291” was still classified as Cluster 2. The reason could be its biased presence in sub-groups. For example, the relative abundance of “476_9291” is higher in samples from China than other two locations. The other two clusters contain OTUs with lower MRA than Cluster 1 and 2, while variations can still be observed within sub-groups.

We further applied a network analysis built by Spearman’s correlations to investigate whether the core OTUs defined were also important to the structure of a co-occurrence pattern. The resulting co-occurrence network contains 293 nodes and 1,424 significant correlations (edges) (Figure S5).

Although most of the core OTUs have relatively low connectivity, they associate with each other rather than with rare OTUs. Nine OTUs were identified as potential “hub” OTUs based on their centrality and the number of links in the network (Figure S5, 6), and as such to community stability. Two core OTUs were identified as hub taxa; “122BU057” (*Megasphaera micronuciformis*) and “524_3631” (*Veillonella atypica*). Compared to other “hub” taxa, they showed lower connectivity and relatively high betweenness centrality (Figure S6).

### 4. Geographic location is the host factor with the largest impact on bacterial composition

To investigate which metadata category has the largest impact on the salivary microbiome, we established random forest models to link the seven categories described above and a new category, study, with the salivary microbiota data at seven taxonomic levels (OTU, phylum, class, order, family, genus, and species). The effect of four normalisation methods was compared using the error rate generated by random forest classification. In total, 224 random forest models were constructed (Figure 5A). Among four normalisation methods, total-sum scaling produced the models that were, on average, the most accurate. Generally, the random forest models built with microbial communities at OTU levels have the lowest error rate (mean = 13.0%), while the models constructed at phylum levels have the highest (mean =26.8%). The model built with the hypervariable region used for sequencing was also the category that showed the lowest error rate (Figure 3G). Geographic locations demonstrated the second important impact on the bacterial communities, with the lowest error rate among biological factors. The random forest model constructed by the other two categories, sample type and tobacco usage, also showed a relatively lower error rate than other categories. Study constructed the models with high error rates at phylum (44.3% ± 2.5%) and class level (29.3% ± 2.1%). However, the error rate of models built with study rapidly dropped with the increase of taxonomic levels, reaching 10.5% ± 5.1% at OTU level. Gender and age range led to poorly performing models.

**Figure 5.**
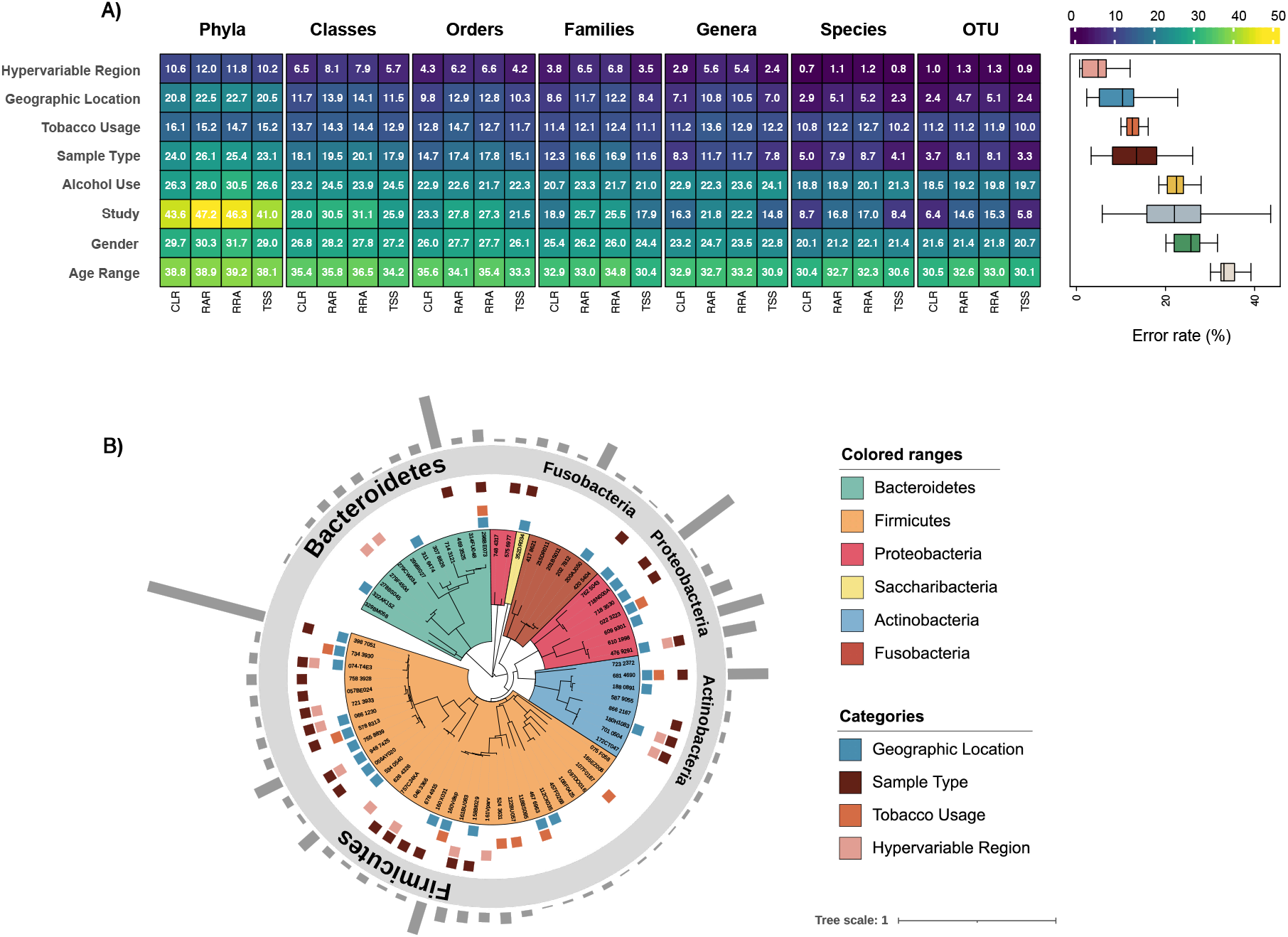
Salivary microbiome members which significantly contribute to categorisation of metadata. Random Forest models showed the impact of categories on salivary microbiome and the core OTUs contributing to accuracy of these models. **A)** Error rate (%) for the random forest classifications conducted with samples grouped by eight different categories. **B)** Phylogenetic tree indicates the taxonomic information of 68 core OTUs. The coloured squares between the tree and the annotation of phylum indicate the OTUs that were defined by the Random Forest model as “important” for distinguishing between different levels in each category. The bars on the outmost ring showing the mean relative abundance of each OTU.

We wanted to determine whether the defined core microbiome could be used as biomarkers to differentiate people categorised by intrinsic and lifestyle factors. The random forest models showed high accuracy at OTU level were used (i.e., geographic location and smoking factors). The differential OTUs induced by hypervariable regions and sample types were also analysed to exclude the influence of technical factors. We further performed ten-fold cross-validation five times to measure the importance of OTUs used to train the model. All OTUs before the point that the cross-validation error curve starts to stabilise were defined as important OTUs. In total, we defined 59, 57, 34 and 70 important OTUs as biomarkers to differentiate samples according to geographic location, smoking habit, hypervariable region, and sample type, respectively (Figure S7). Of these, 28, 10, 13 and 22 biomarkers were also classified as “core” (Figure 5B; Figure S8). Although 31 core OTUs showed the importance in discriminating samples according to geographic location and smoking, nearly half of them (15 OTUs) had the possibility of being confounded by technical factors (Figure 5B). After excluding the OTUs that could be influenced by other factors, core OTU “322AK152” (*Bergeyella sp*.*HMT_322*) was the OTU with the highest contribution to the classification of samples from three geographic locations. Meanwhile, “122BU057” (*Megasphaera micronuciformis*) showed the highest importance among the core differential OTUs specific to smoking, followed by “524_3631” (*Veillonella atypica*). We were surprised to find that these two OTUs were the only two OTUs that were defined as hub taxa in the co-occurrence network analysis (Figure S6).

### 5. The salivary microbiota as biomarkers to differentiate Chinese and Western participants

We further analysed the changes caused by geographic locations in higher taxonomic hierarchies, where many differences have been revealed. Of particular interest were taxa under phylum *Synergistetes* and *Spirochaetes*, Class *Mollicutes* and *Betaproteobacteria*, Family *Clostridiales*, and genus *Prevotella*. Interestingly, many taxa showed a higher relative abundance in the Chinese samples, both compared to North American samples and compared to European samples (Figure 6A). It suggested that the variance induced by geographic locations may be dominated by the differences between samples from Chinese and Western people. Therefore, we combined samples from North America and Europe into a single group, “Western.” Compared to the Chinese grouping, the Western group has significantly lower within-sample diversity (alpha-diversity) (Wilcoxon rank-sum test, p < 0.001; Figure 6B, C). Next, we examined the differences between Chinese and Western in the salivary microbiota at the genus and species level (Table S6, S7). Besides establishing a random forest model, we also identified differential taxa using ANCOM-BC, adjusting for the hypervariable region. We found 48 genera identified as significantly different by both methods (Figure 6D, Table S6).

**Figure 6.**
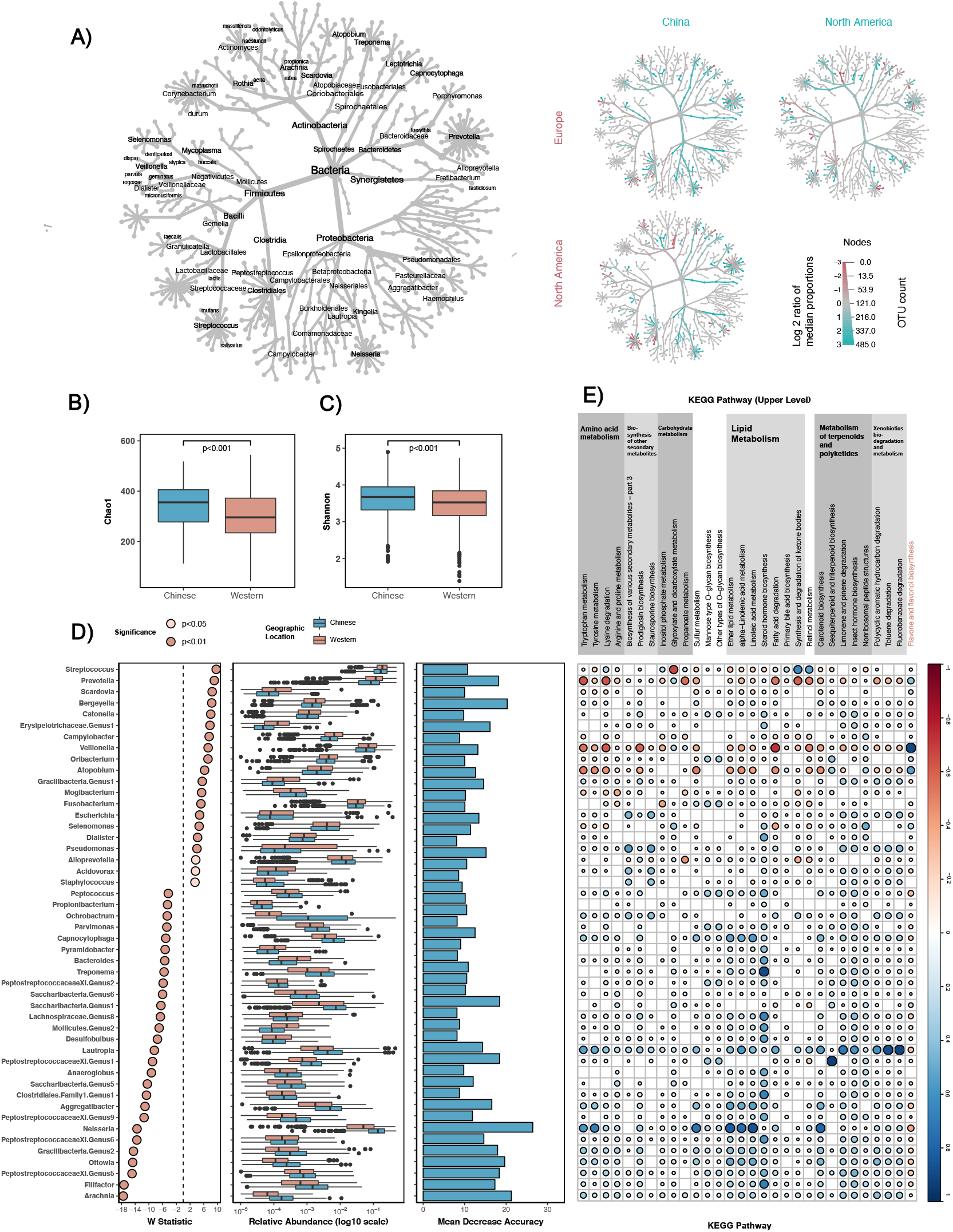
Distinct microbial profiles are evident in the saliva samples from Chinese and Western adults. **A)** Taxonomic hierarchies show the relative enrichment of taxa in three geographic locations at phylum through species level. Coloured nodes indicate log2-fold increase in median abundance of the group in x-axis (pink) or y-axis (blue). Only taxa showed significant changes (false discovery rate-adjusted Wilcoxon rank sum q < 0.05) are displayed. **B)** and **C)** Comparison of salivary microbial alpha diversity between the Chinese and Western samples, calculated by Chao1 (B: *p*< 0.001, Wilcoxon rank-sum test) and Shannon index (C: *p*< 0.001, Wilcoxon rank-sum test). **D)** Differential abundant genera identified between saliva from Chinese and Western samples. The panel on the left indicates the standardised effect sizes (W statistic) estimated via the difference on relative abundance using ANCOM-BC (taxa enriched in Western samples have a value shifted to right, whereas taxa enriched in Chinese samples have a value shifted to left); The panel in the middle shows the relative abundance of selected genera; the panel on the right indicates the Mean Decrease Accuracy of the random forest model established. **E)** Spearman’s correlation coefficients were calculated between each pairwise comparison of differential genus and KEGG pathway. Only significantly correlated comparisons (p<0.01, FDR adjusted Spearman’s rank correlation) are displayed. The only Western-enriched pathway is marked in pink.

Finally, we performed the functional prediction-based 16S rRNA gene profiles to investigate whether differences in the salivary microbiota between Chinese and Western affect its function. Two methods, ANCOM-BC, and random forest model were used to identify which pathways were differential between Chinese and Western. The result of ANCOM-BC indicated that 69 pathways related to metabolism were differentially abundant between the two groups. The random forest classification model established using KEGG pathways demonstrated an error rate of 10.01% and revealed 46 differential pathways. Among them, thirty pathways belonging to nine upper pathways (level 2) were simultaneously defined by two methods as differing in abundance between Chinese and Western (Figure 6E, Table S8). A variety of pathways was in higher abundance in Chinese samples. The enrichment of these pathways in Chinese samples was mainly associated with the increased abundance of *Neisseria* and *Lautropia* and the depleted abundance of *Prevotella, Veillonella*, and *Atopobium*. Notably, three lipid metabolism pathways enriched in Chinese samples, including “Ether lipid metabolism” (ko00565), “alpha-Linolenic acid metabolism” (ko00592), and “Linoleic acid metabolism” (ko00591), have the highest standardised effect size (W statistics, Table S8). The enrichment of these pathways related to lipid metabolism has been positively associated with the higher abundance of *Neisseri*a in Chinese. *Neisseria* may have also contributed to the pathway “Carotenoid biosynthesis” (ko00906). Another metabolic pathway related to the metabolism of terpenoids and polyketides, “Sesquiterpenoid and triterpenoid biosynthesis” (ko00909), showed a positive correlation with a genus belong to *Peptostreptococcaceae*. In contrast, only one pathway named “Flavone and flavonol biosynthesis” (ko00944) was enriched in the saliva samples from Western. A strong positive correlation has been demonstrated between this pathway and the increased abundance of *Veillonella* in the samples from the Western grouping.

### 6. Validation of the core in an independent Australian cohort

To validate the prevalence of the core OTUs in human saliva, we collected saliva samples from 13 Chinese and 13 Western participants in Melbourne and sequenced the extracted DNA with 515F-806R primers. In total, 841,188 high-quality 16S rRNA sequences were obtained, which clustered into 397 OTUs with 97% identity to the HOMD database. Among them, the core OTUs we defined in the meta-analysis showed high relative abundance (78.3 ± 6.9%) in all collected samples. To increase the accuracy of the OTU assignment, we denoised the sequencing reads using the UNOISE3 pipeline and generated ZOTUs with 100% sequence identity. After re-clustering the core OTUs defined in the meta-analysis to ZOTUs, we observed that 59 of the identified OTUs in this independent dataset consisted of 87 ZOTUs, and thus made up close to 80% of the relative abundance (Figure 7A). Although some sequences belonging to the same ZOTU are clustered to different OTUs, all the core OTUs contain at least one highly abundant ZOTU. The taxonomic profiles of the global core annotated by the HOMD database and the ZOTUs annotated by the SILVA database were very similar at the genus level (Figure 7A).

**Figure 7.**
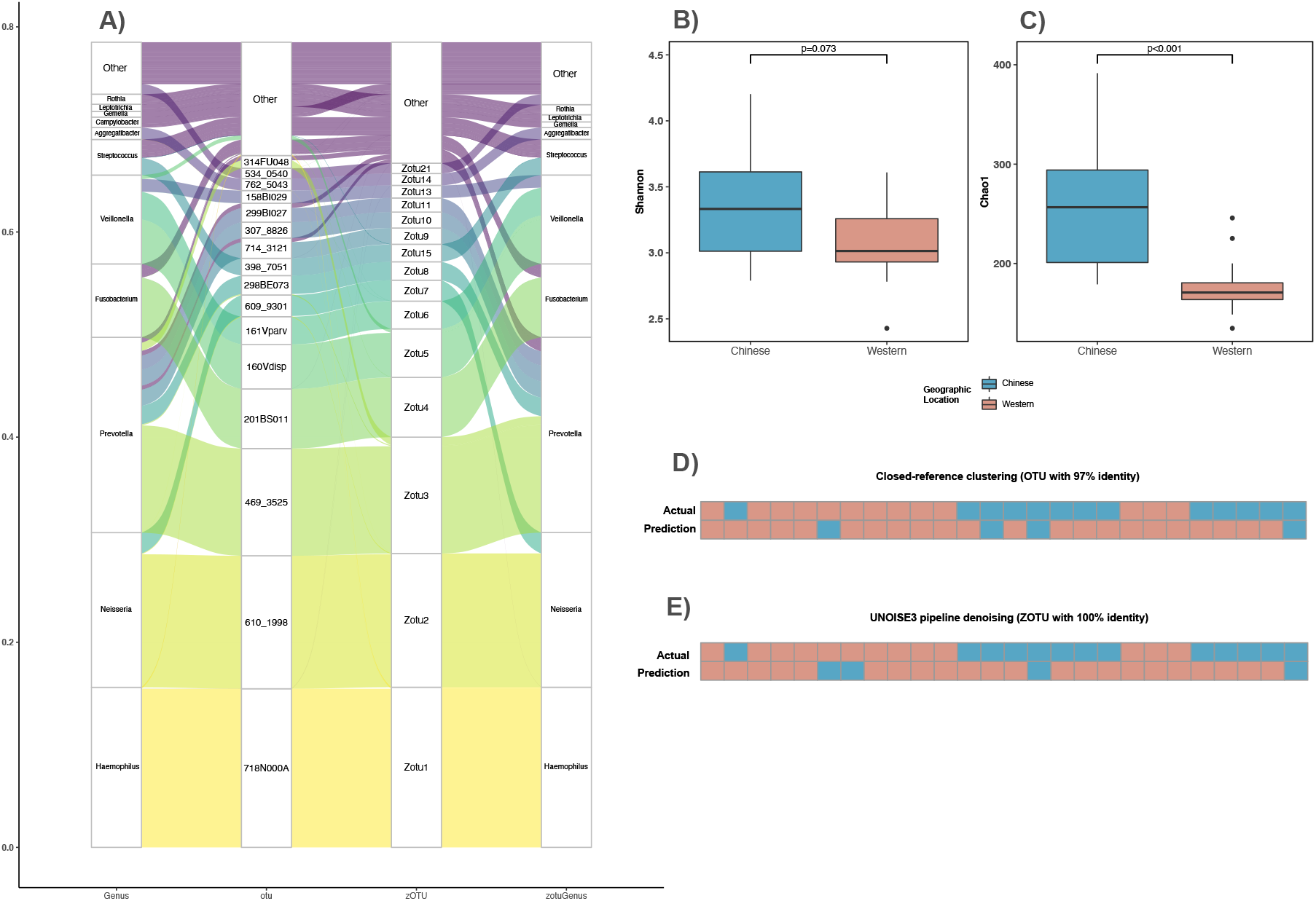
An independent cohort verifies the definition of core microbiome membership but cannot classify based on cultural background. **A)** Alluvial plot showing the affiliation of ZOTUs to their originating core OTUs defined in the meta-analysis. **B)** and **C)** Comparison of salivary microbial alpha diversity between the Chinese and Western samples, calculated by Shannon (B: *p* = 0.073, Wilcoxon rank-sum test) and Chao1 index (C: *p*< 0.001, Wilcoxon rank-sum test). **D)** and **E)** The prediction of the cultural backgrounds of the samples according to the random forest classification model constructed using the genus profiles of samples in the meta-analysis. The genus level profiles of samples processed by **D)** closed-reference clustering with 97% sequence identity and **E)** UNOISE3 denoising with 100% sequence identity were used as the test set.

We wanted to verify the observed differences between the OTUs in saliva samples from Chinese and Western people in this independent dataset. Although two groups did not differ significantly when considering the Shannon diversity index (Wilcoxon rank-sum test, p = 0.073; Figure 7B), the Chinese group showed a higher Chao 1 Index than the Western group (Wilcoxon rank-sum test, p < 0.001; Figure 7C), which is in agreeance with the result of the meta-analysis. Meanwhile, no significant differences were observed between the two groups by beta-diversity analyses (Bray-Curtis and weighted uniFrac distance, Table S9). We used the random forest classification model constructed using the genus level profile of the large-scale dataset to predict the Chinese and Western samples in this independent study. This dataset’s genus-level relative abundance table was prepared from both the OTU table with 97% identity to HOMD v15.1 database (Figure 7D) and the ZOTU table annotated by the HOMD v15.22 database (Figure 7E). The accuracy of both predictions was relatively low, with 57.7% for the OTU table and 50% for the ZOTU table. Interestingly, most Western samples were correctly classified, while most samples from Chinese participants were classified as being ‘Western’ in this analysis.

## Discussion

There is ample knowledge on the disease-affected salivary microbiota, yet our perspectives to the bacteria present in healthy humans remains limited. Our systematic selection of studies, together with the re-analysis of the 16S rRNA amplicon sequencing data from 47 studies offers a comprehensive description of the salivary microbiome presented in adults without diagnosed disease. Our study has defined the core members of salivary bacterial communities across 2211 samples from 47 studies and has used metadata captured in these studies to investigate the role of different intrinsic and extrinsic factors on the occurrence of these core. It is clear that core members differ between geographic locations of collected saliva, and our analysis shows that Chinese participants are different from Western participants (encompassing European and North American studies). A prediction of the pathways enriched in each collective indicates that bacterial metabolic pathways are likely to influence the aroma and flavour perception of foods. These results show that despite the core microbial members of saliva being common across humans, there are differences, likely due to diet. We suggest that the aroma and flavour of foods and beverages are likely to be differently affected in healthy humans across the globe, meaning that preference and consumption of different foods is likely to be prioritised. These results have important consequences for food and beverage design, composition, and dietary advice across the globe.

Based on the abundance-occupancy pattern, the definition of core microbiome highlighted the persistent and conserved microbial communities in human saliva across the globe. Here, we compared two approaches adapted from two studies (references) to defining the core. The method adapted from the study of Wu et al. [13] is a relatively conventional strategy that has been chosen by many other studies with customised thresholds [45-47]. For another approach we applied in this study [35], we determined the occupancy of OTUs according to their detection over study. When evaluating the contribution of top-ranked OTUs in occupancy to the beta-diversity of the community, we further used weighted Unifrac due to its effectiveness in minimising the biases induced by the selection of hypervariable regions.

The resulting core members identified by these methods have a lot in common. A majority of the core OTUs defined by Shade’s method is also included in the cores defined by Wu’s method, suggesting the recently developed multi-step approach is effective in determining taxa with high prevalence. The general high relative abundance of the core across different sub-groups emphasised the utility of this pipeline in identifying the persistent members across diverse datasets. Most of the core salivary microbiota we defined had been proposed in previous studies as prevalent bacteria in the human oral cavity that persistently span across different individuals [11, 48-50]. The dominant genus of the core we defined, *Streptococcus, Neisseria*, and *Prevotella*, were concluded as core human salivary microbiome by a recent study based on the MG-RAST data [51]. Ten OTUs belonging to genus *Streptococcus* were included in the “core”, two of which were classified to the cluster with overall the highest relative abundance across all dimensions. The prevalence of *Streptococcus* we observed is consistent with a previous study defining the healthy core from the 454 pyrosequencing results of three individuals [11]. The most abundant core OTU we found, *Streptococcus oralis* subspecies *dentisani*, has been documented in previous studies as potential oral health-promoting organisms and being highly abundant at various oral niches of healthy humans [52, 53].

We further conducted a co-occurrence network analysis to investigate the role of these core microbiota in shaping the microbial community and found the co-existence between many members of the core (Figure S5). The presence of the rare OTU that became the hub suggests that although some taxa are not persistently detected across the community, they may still be important for the overall structure of the salivary microbiome. It has been proposed that the oral microbiota of healthy individuals is both homeostatic and dynamic [48]. The core microbial members consistently present in human saliva identified here may explain the stability of oral microbiota to some extent. Moreover, we conducted an independent study to verify the prevalence of the core defined from the published studies. The high relative abundance and occurrence of the original core OTUs in this independent cohort suggest that the core human salivary microbiome we defined can be applied to different datasets. After re-clustering the core OTUs to ZOTUs with 100% sequence identity, we may conclude that the same members constitute the core, even if different taxonomic resolution is applied.

Besides the core microbiome, there are “variable” microbiota in human microbial communities, which vary among individuals because of unique lifestyle and genetic factors [54]. We performed analyses for beta-diversity of samples (Figure 3) and random forest classifications (Figure 5A), demonstrating several factors-both technical and physiological-significantly discriminated between sub-populations. Because of the high heterogeneity between studies in their methodology, large inter-study variability was the main factor that affected the observed salivary microbiota [55, 56]. For example, the impact of chosen hypervariable regions for sequencing on driving microbial community structures has been confirmed by our study. Inter-study variation may also be attributed to the criteria of recruiting participants. Although all samples included in this meta-analysis were collected from the control groups and population without specific diagnosed disease, the definition of ‘healthy’ varies. For example, the use of antibiotics was not always considered as an exclusion criterion, and when included, different time intervals were adopted. However, we aimed to construct a microbial community that reflects the salivary microbiota of real-life consumers.

The importance of various host intrinsic and lifestyle factors, especially smoking, changed the salivary microbial composition [57]. We identified a core OTU belonging to *Megasphaera micronuciformis* as a biomarker for reported smoking [58, 59]. *Veillonella atypica* OTU’s vary between smokers and non-smokers, agreeing with a previous study [60]. Another strong determinant of the salivary microbiome we defined was “sample type”. As we found in this study, the difference between the mouthwash sample and the other two collection methods is greater than the difference between the stimulated and unstimulated saliva. Contrary to the result of Jo et al. [29], the OTU belonging to *Neisseria flava* has not been identified as differential taxa for the type of saliva. The microbial composition of alcohol drinkers was different to non-drinkers, but small sample size hinders other conclusions.

Geographic location has been identified as the host physiological factor with the largest impact on salivary microbiota (Figure 5A). Although it only explained limited variability between samples’ microbial profiles, the observed variations were robust to the heterogeneity induced by different hypervariable regions used (Figure 3E, F). To date, little is known regarding the influence of geographic locations on the human salivary microbiome. A comparative study reported the differences in saliva microbial composition in Alaskans, Africans, and Germans [33]. Our result extends this study to demonstrate that core OTUs may differentiate saliva samples from North America, Europe, and China. Given the high abundance and occupancy frequency of the core microbiome, we would expect these taxa to be effective indicators to predict the geographic background of saliva donors.

Due to the sometimes large differences in culture and lifestyle between Western and non-Western populations, we further grouped our data into Western and Chinese samples. The comparison between Western and non-Western populations has already been applied to the human gut microbiota, whereas less is known about the saliva microbiota [61]. Our study found the difference between saliva fromcc Western and Chinese people in the abundance of *Veillonella* spp. Here, we showed that *Veillonella* was generally higher in Western samples. Such differences may further influence the flavone and flavonol biosynthesis pathways in the oral cavity. Our previous study revealed the Western-born and Chinese-born wine experts had different responses to the flavour of wine (unpublished results: 10.13140/RG.2.2.11263.12965) [62]. Since flavonol is a well-known constituent of wine-related to the bitterness and astringency perception [63], we would hypothesis that the enrichment of *Veillonella* in Western may affect their sensitivity to the phenolic compounds in wine. Regular consumption of flavonoid-rich foods, such as oolong tea, may increase the abundance of *Veillonella* spp. in human saliva [64]. These could link the differences between Chinese and Western groups in sensory evaluation and their salivary microbiota together.

Although the independent dataset we collected did not show differences between samples taken from Chinese and Western participants, the results of random forest classification may lead to some interesting hypotheses. The prediction of models revealed that most of the Chinese samples in this cohort were classified as Western. The donors of these samples were wine experts, of Chinese ethnicity, born in China and living in Australia for no more than 18 months. Recent studies reported that immigrants from Asia experience a “Westernization” of gut microbiota induced by dietary acculturation [65, 66]. We hypothesise that such a phenomenon may also happen in salivary microbiota. It may suggest that dietary pattern is a more important determinant than ethnicity in shaping the salivary microbiota of the participants, leading to variation among different geographic locations that we observed in the meta-analysis. In summary, we have defined a core bacterial community in saliva from healthy humans, and this core demonstrated both stability and variability among populations. The prevalence of the core members of the saliva microbiome has been confirmed in an independent cohort. We have revealed the influence of various host factors, such as geographic locations, incidence of smoking and drinking, on the salivary microbiome. We also identified microbial and functional biomarkers to differentiate the Chinese and Western people, underlying the potential relationship between salivary microbiota and sensory perception. Results in this work will provide foundational information to inform future studies to understand the similarities and differences in saliva microbial composition, potentially associating oral to aroma and flavour perception of foods.

## Declarations

### Ethics approval and consent to participate

The study was approved by the Office for Research Ethics and Integrity of the University of Melbourne (Ethics ID: 1852616), and all participants provided written informed consent.

### Data availability

The sequencing data supporting the conclusion of the meta-analysis in this article are available in publicly accessible databases (full details can be found in Table S2). The sequencing data generated and/or analysed during the current study are available in the NCBI Bioproject repository, PRJNA786805 (https://www.ncbi.nlm.nih.gov/bioproject/786805). Original scripts generated during the current study are available in Github (https://github.com/XINWEIR/SalivaryMicrobiome_MetaAnalysis).

### Authors’ contributions

The research topic was developed by XR and KH. XR conducted the meta-analysis with the guidance of KH. XR and JL collected and analysed the local samples. XR wrote the first draft of the manuscript and all authors contributed to revise it. All authors read and approved the final manuscript for submission.

## Acknowledgements

This study was funded by the Faculty of Veterinary and Agricultural Sciences at the University of Melbourne. JL and XR gratefully acknowledge a Melbourne Research Scholarship administered by the University of Melbourne.

