## Supplementary material for "The salivary microbiome shows a high prevalence of core bacterial members yet variability across human populations": Supplementary Figures.docx

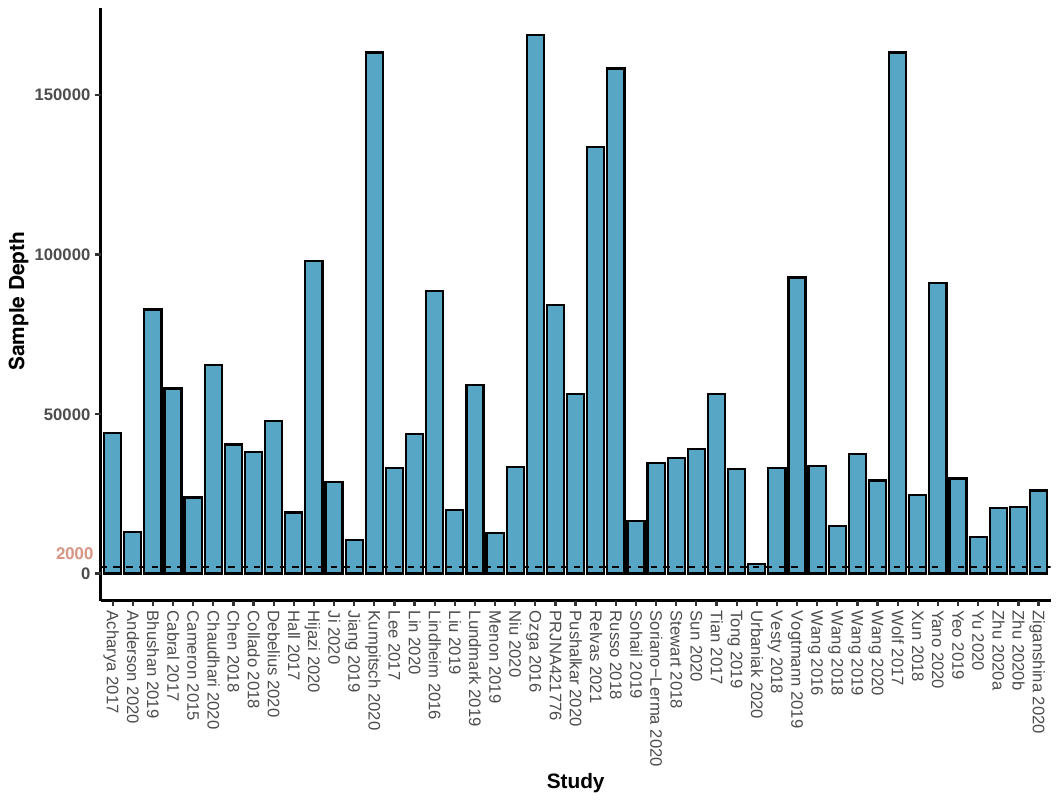

**B)**

**A)**

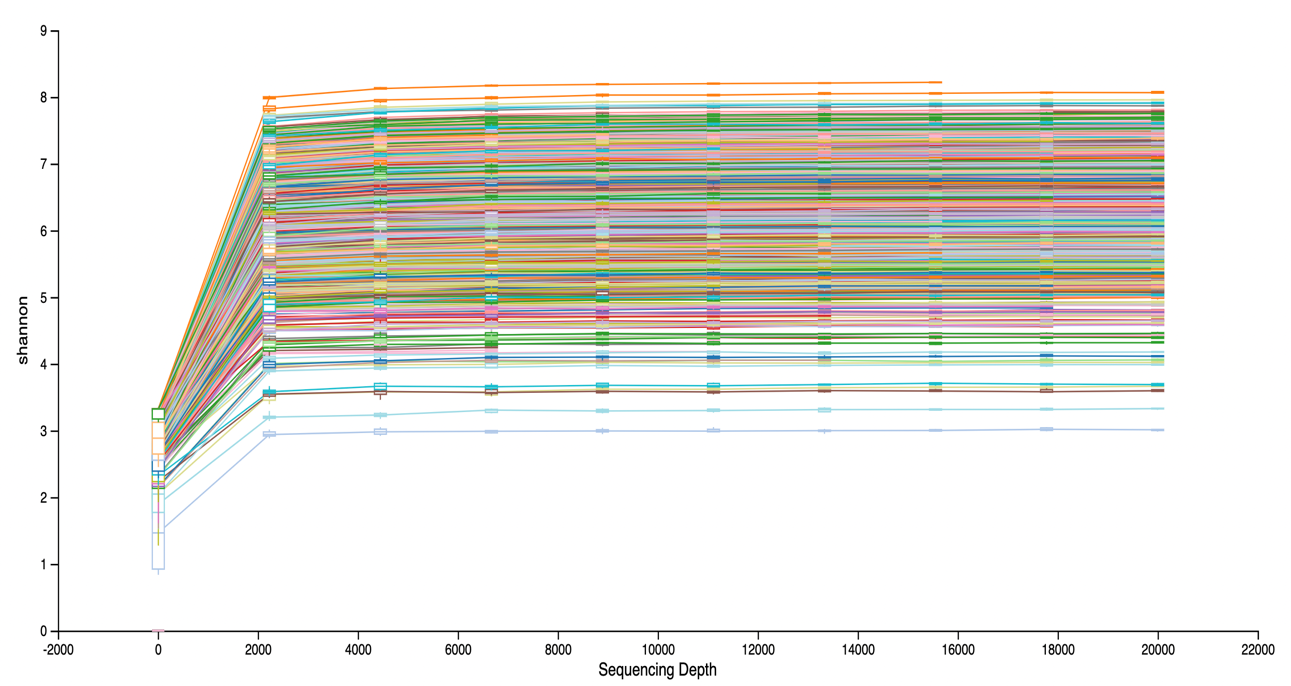

**Figure S1 Average sequencing Depth and rarefaction curve for the whole 16S rRNA dataset. A)** Mean read number of samples from each study. The dash line indicates that all samples below this depth (depth = 2,000) have been removed. **B)** The rarefaction curve reflects the increase of sample’s Shannon index with sequencing depth. The curve was basically stabilised at sequencing depth = 2000.

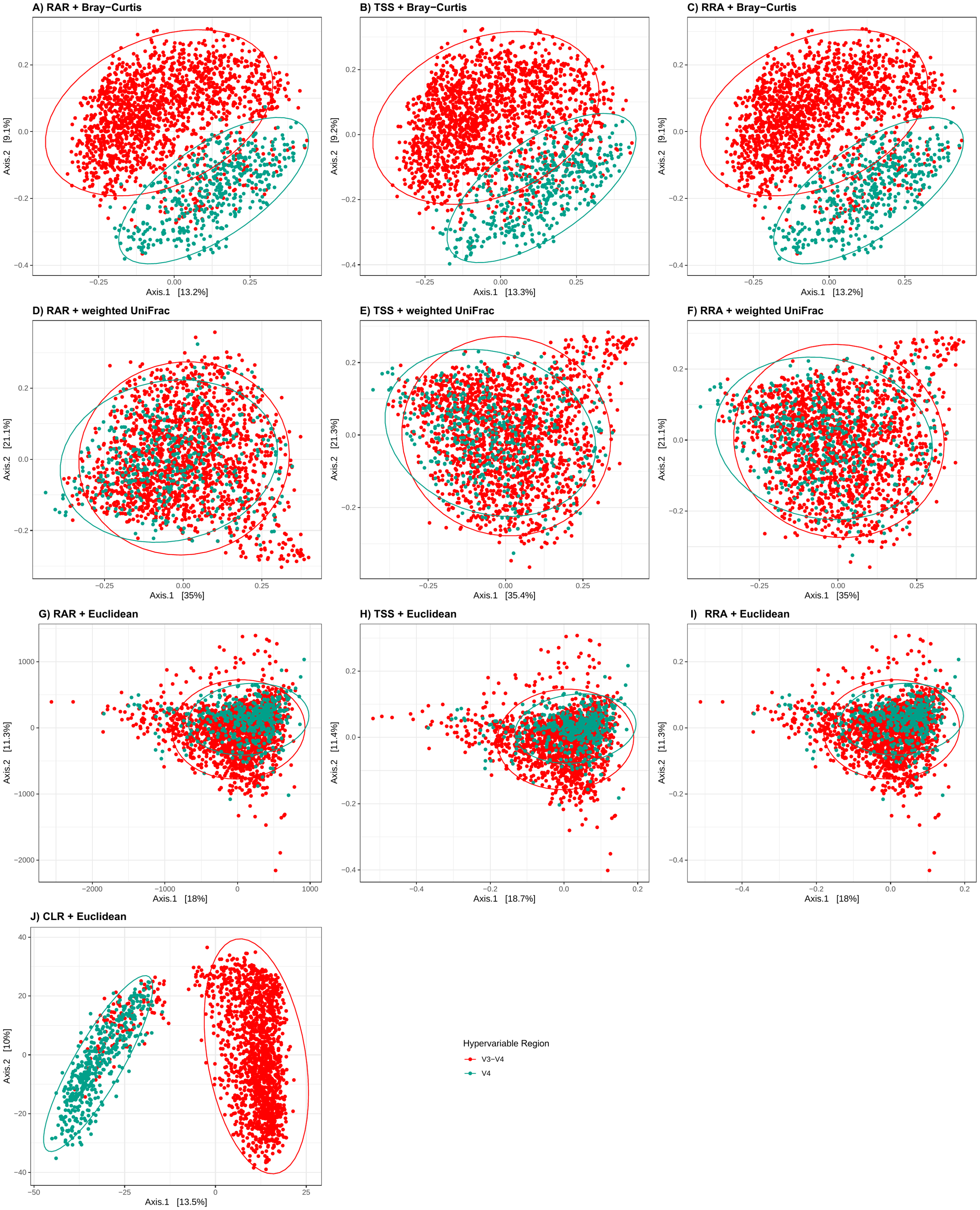

**Figure S2 The effect of different combinations between normalisations and distance matrices on reducing the impact of hypervariable regions in the PCoA plot.** PCoA plots showing Bray-Curtis dissimilarity **(A-C)**, weighted uniFrac distance **(D-F)**, Euclidean distance **(G-J)** under rarefaction (RAR) **(A, D, G)**, total-sum scaling (TSS) **(B, E, H)**, rarefied relative abundance transformation (RRA) **(C, F, I)** and centred log-ratio transformation **(J)**. Percentage of variances explained by the first two principal coordinates are shown on the axes.

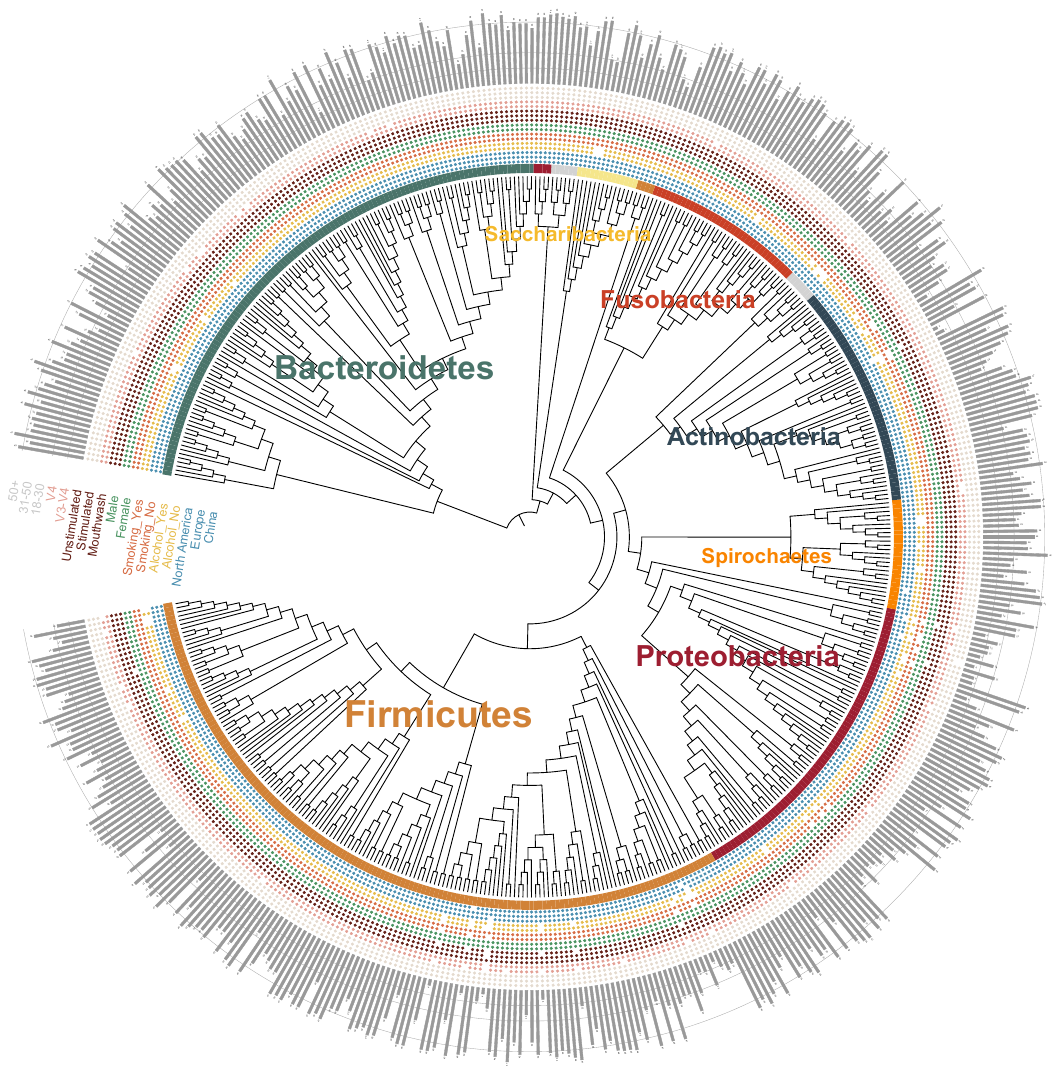

**Figure S3 Phylogenetic tree showing the presence and absence of top 500 OTUs with highest mean relative abundance.** The colour strips on the innermost ring indicates which phylum the OTUs belong to. The presence of coloured circles on the 17 rings in the middle indicate that an OTU was found in a specific level of a sub-group. The grey bars on the outermost layer represent how many studies that an OTU presented. The scale lines are used to highlight the number of 10, 20 and 40.

.

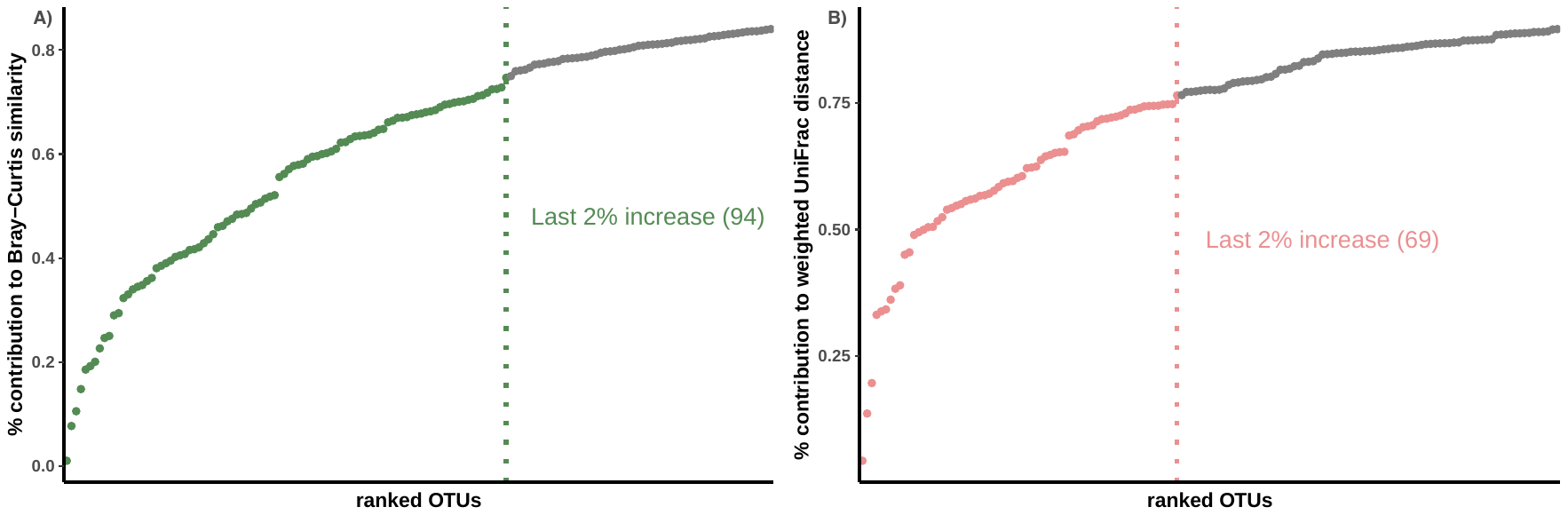

**Figure S4 The percentage contribution of the top-ranked OTUs to the beta-diversity of the dataset.** The beta-diversity is calculated for the whole dataset and for only the top-ranked OTUs using both Bray-Curtis similarity (A) and weighted uniFrac distance (B). The percentage contribution of top-ranked OTUs is calculated by dividing the beta-diversity among top-ranked OTUs using the beta-diversity of the whole dataset. The dash lines indicate the last points at which the increase on the contribution is 2%. All OTUs before this point (the point itself was also included) were defined as “core”.

**Figure S5 Bacterial co-occurrence network verifies the role of identified core salivary microbiome members.** Small modules with less than seven nodes were not displayed. The size of nodes is proportioned to the connectivity of nodes (node degree). Core OTUs from Figure 4 are indicated as yellow, while rare OTUs are in green. The edges between nodes represent the strength of the correlation (Spearman’s correlation coefficient, ρ ≥ 0.5, p < 0.01). The shape of “hub” OTUs are indicated as squares with the OTU name displayed.

**B)**

**A)**

**Figure S6 The Betweenness and Closeness centrality of OTUs involved in the network analysis.** The “hub” OTUs were identified as OTUs with either high connectivity (node degree) or centrality (betweenness **(A)** and closeness **(B)** centrality). The accession number of “hub” OTUs are indicated. The core OTUs are marked in yellow, while the rare OTUs were coloured in green.

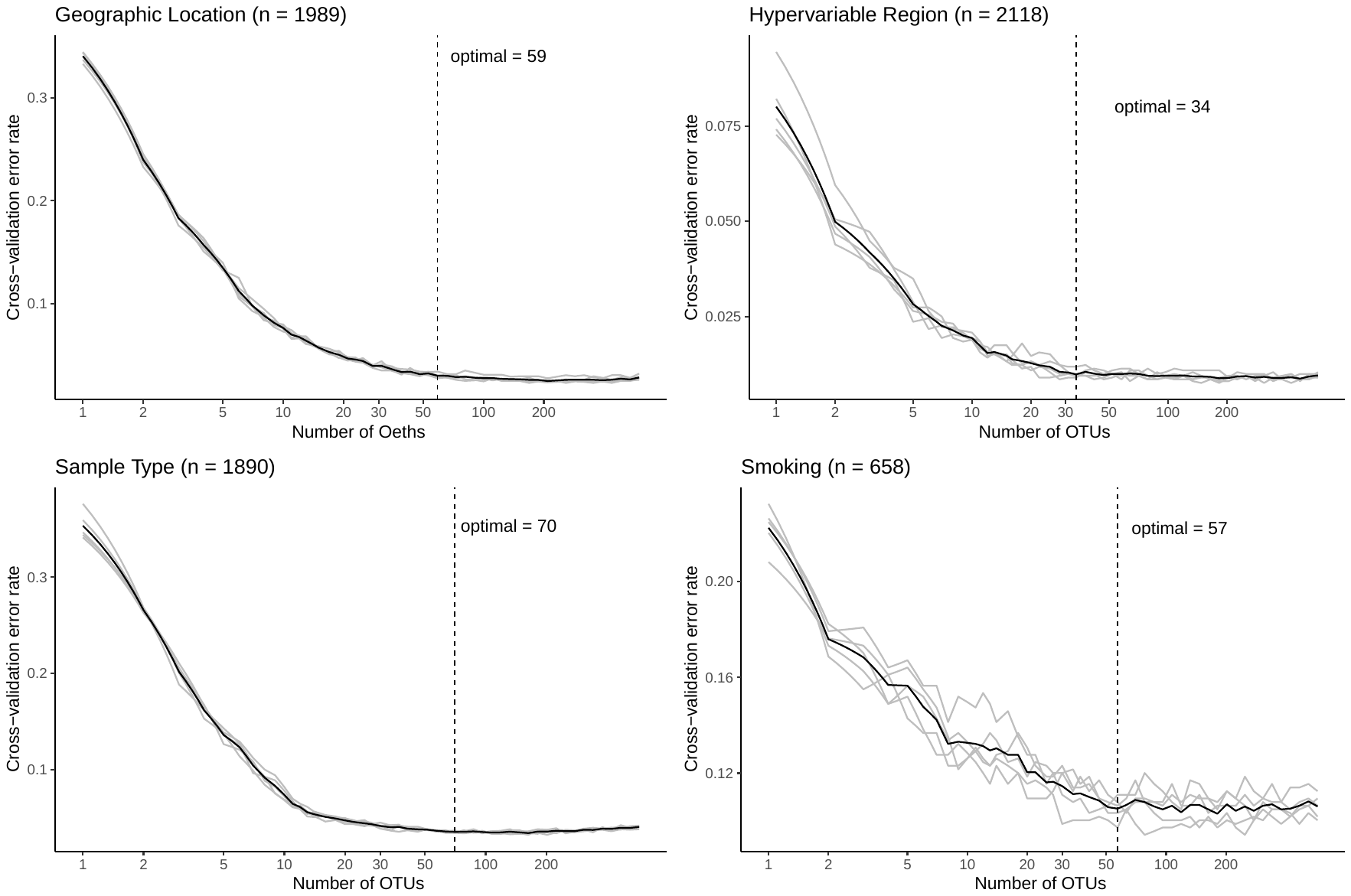

**Figure S7** **The optimal number for defining the biomarker OTUs of four categories.** The contribution of the OTUs used to differentiate the levels in each category on ten-fold cross-validation. The OTUs were ranked in the order of importance. The dash lines represent the point at which the curve starts to become overall stable.

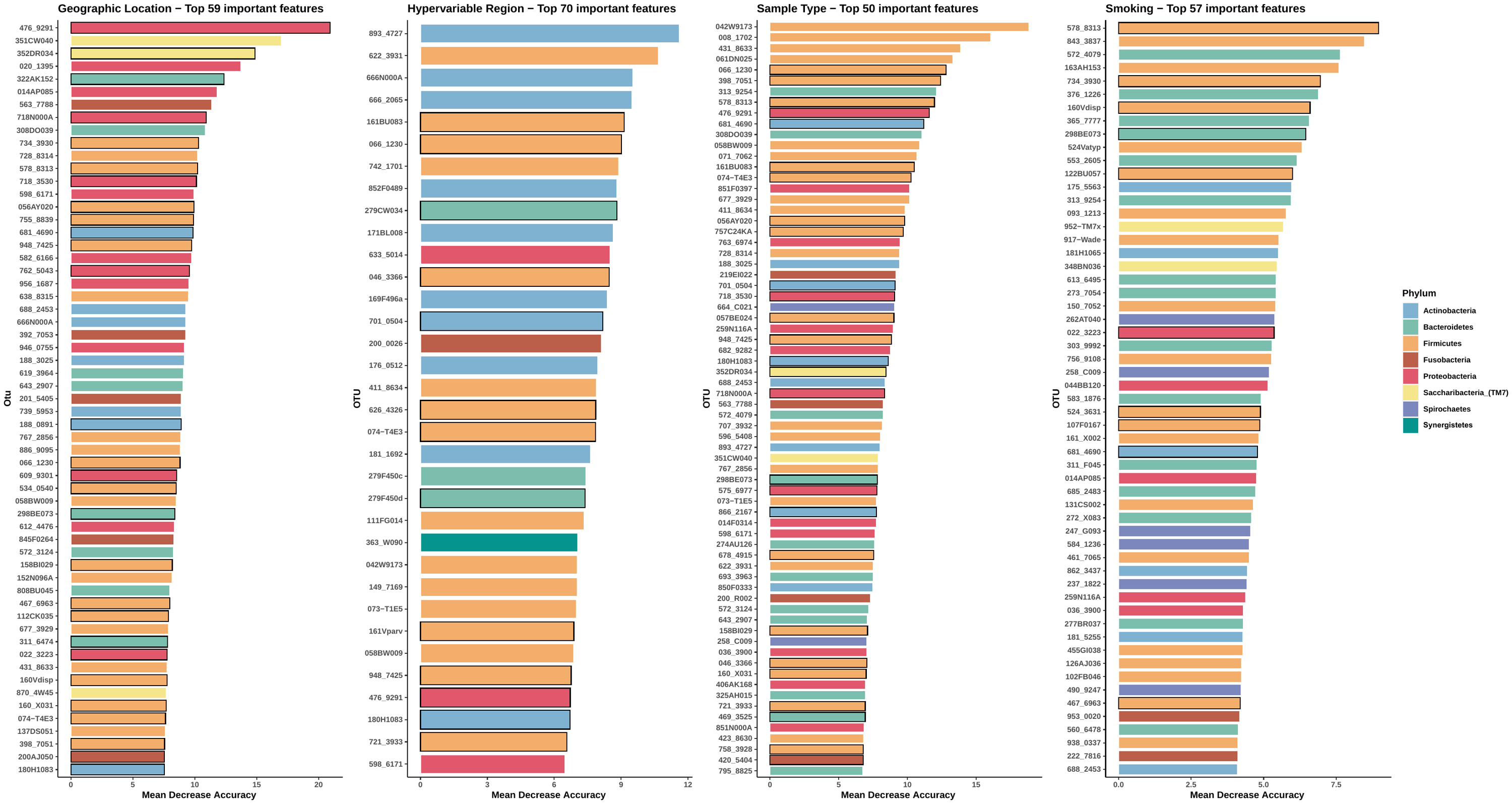

**Figure S8 The top important OTUs identified by Random Forest classification model established by the relative abundance of human salivary microbiota.** The differential OTUs defined were ranked in descending order of their importance (Mean Decrease Accuracy). The colour of bars reflects the phylum level information of OTUs. The Mean Decrease Accuracy bar of core OTUs were marked with bolded black borders.
