## Supplementary material for "The salivary microbiome shows a high prevalence of core bacterial members yet variability across human populations": TableS1_Searching_Terms.docx

| **TERMS** | **Medline and EMBASE** | | **Web of Science** |
| --- | --- | --- | --- |
| **Saliva or Oral** | **Map Term to Subject Heading** | Saliva/ | Saliva* or oral near/5 (microbiome OR microbiota OR bacteria*) |
|  | OR | |  |
|  | Keywords | Saliva* or oral adj5((microbe* or microbiome* or microbiota* or bacteria* or microbial) or Microbiota/ or exp Bacteria/) |  |
| AND | | | |
| **Microbiome** | **Map Term to Subject Heading** | Microbiota/ or exp Bacteria/ | Microbe* OR microbiome* OR microbiota* OR bacteria* OR microbial |
|  | OR | |  |
|  | Keywords | microbe* or microbiome* or microbiota* or bacteria* or microbial |  |
| AND | | | |
| **16S rRNA amplicon sequencing** | **Map Term to Subject Heading** | RNA, Ribosomal, 16S/ | “RNA, Ribosomal, 16S” OR “RNA 16S” OR 16S |
|  | OR | |  |
|  | Keywords | 16S |  |

**Table S1 Searching Terms used on Medline, EMBASE, and Web of Science.**
